# A Pan-Cancer Benchmark of Clustering Strategies for Patient-Specific Gene Regulatory Networks

**DOI:** 10.64898/2026.10.20.759648

**Authors:** Panos Sarantidis, Tatiana Belova, Mariike L. Kuijjer

## Abstract

Cancer patients with clinically similar tumors can exhibit substantial differences in disease progression and treatment response. Patient-specific gene regulatory networks (GRNs) may capture regulatory heterogeneity that is not represented by gene expression alone. However, strategies for clustering patients using these high-dimensional networks have not been systematically evaluated and it remains unclear whether clustering based directly on millions of patient-specific regulatory edge weights is computationally feasible at scale. We benchmarked feature-selection and clustering approaches for patient-specific GRNs inferred with PANDA and LIONESS across 9,778 primary tumor samples from 33 cancer types. We compared three data representations: gene expression, gene indegrees summarizing regulatory input to each gene, and complete networks of transcription factor–gene edge weights. Twenty combinations of four feature-selection methods and five clustering algorithms were evaluated using consensus clustering. Performance was assessed based on mean silhouette score, PAC score, concordance, computational time, and memory usage. Standard deviation–based feature selection combined with k-means clustering achieved the strongest overall performance for both GRN-derived representations, with overall scores of 0.997 for edge weights and 0.985 for indegrees, while remaining computationally efficient. Edge-weight- and indegree-derived clusters showed clinical, biological, and prognostic associations that were complementary to those obtained from gene expression. Together, these findings provide a practical framework for network-based cancer patient stratification and highlight the potential of GRN-derived features to reveal clinically relevant tumor heterogeneity beyond gene expression alone.

## Introduction

Cancer is a highly heterogeneous disease, and patients diagnosed with clinically similar malignancies frequently display substantial variation in disease progression and therapeutic response. Advances in molecular profiling technologies have enabled increasingly detailed characterization of tumor heterogeneity. Clustering approaches applied to molecular data can group tumors with similar molecular profiles and have therefore become widely used for cancer subtyping, with patients assigned to the same cluster often sharing similar clinical characteristics (Jamalinia and Weiskirchen 2025). Gene expression data have been extensively used for this purpose (Oyelade et al. 2016). However, expression-based analyses do not explicitly model the regulatory interactions between transcription factors and genes that may drive tumor heterogeneity.

Gene regulation is widely recognized as a core driver of numerous biological processes and is thought to play an important role in cancer development and progression (Lee and Young 2013; Bradner et al. 2017). Transcriptional regulation is mediated through interactions between transcription factors and their target genes, which together form regulatory programs associated with cellular state. Gene regulatory networks (GRNs) provide a framework for modeling these interactions. They have been widely used to investigate cancer-associated regulatory programs, identify dysregulated pathways, and characterize genes involved in tumor progression (Carson et al. 2015).

Several computational methods have been developed to infer population-level GRNs from transcriptomic data. Among these, PANDA integrates prior regulatory information, protein–protein interactions, and gene co-expression data to reconstruct complete, weighted networks, in which edge weights represent regulatory interaction likelihoods estimated across a population of samples (Glass et al. 2013). However, population-level networks do not capture patient-specific regulatory variation. LIONESS is a single-sample network inference methodology that can be applied together with PANDA to estimate GRNs for individual patients (Kuijjer et al. 2019), enabling direct characterization of patient-level regulatory variation and supporting downstream analyses focused on inter-patient heterogeneity (De Marzio et al. 2023; Belova et al. 2025).

Although patient-specific GRNs are increasingly used in cancer studies (Saha et al. 2025; Deschildre et al. 2024; Lopes-Ramos et al. 2021), their use for clustering-based patient stratification has not been systematically investigated. Clustering of patient-specific GRNs has so far been mainly performed on summarized network representations, such as node degrees or pathway-derived features, rather than directly using the complete regulatory interaction structure. One commonly used summarized representation is the gene indegree, which captures the total regulatory input received by a gene across all incoming transcription factor interactions, thereby substantially reducing dimensionality (Weighill et al. 2021). In contrast, edge-weight representations preserve the complete transcription factor–gene interaction landscape. However, clustering directly on these full edge-weight representations remains largely unexplored because of the extreme dimensionality and the computational demands associated with analyzing these data. This represents an important computational challenge for network-based patient stratification, particularly for large patient cohorts.

In addition to the choice of network representation, the identification of robust patient subgroups using patient-specific GRNs depends on the applied clustering strategy. High-dimensional molecular datasets contain large numbers of features that may obscure biologically meaningful structure and reduce clustering robustness. Feature-selection approaches are therefore commonly used to prioritize informative variation before clustering. Moreover, clustering algorithms may produce substantially distinct subgroup structures because of differences in their underlying assumptions and optimization procedures. Despite the importance of these methodological choices, systematic benchmarks of clustering strategies for patient-specific GRNs have, to our knowledge, not yet been reported, particularly for high-dimensional edge-weight representations.

In this study, we benchmark clustering strategies for patient-specific GRNs inferred using PANDA and LIONESS across 33 cancer types from The Cancer Genome Atlas (TCGA) (The Cancer Genome Atlas Research Network et al. 2013). We compare clustering based on gene expression, gene indegrees, and full edge-weight representations, and systematically evaluate combinations of feature-selection and clustering approaches with respect to computational load, clustering robustness and biological relevance, with the goal of identifying methods that produce robust, high-quality clusters that could support our understanding of cancer heterogeneity. In particular, we evaluate whether edge-level clustering is feasible at scale and identify strategies that make such analyses practical across diverse cancer types. We further assess associations between identified clusters and clinical variables, pathway enrichment, and patient survival, and find that GRN-derived representations can capture biologically and clinically relevant information beyond that captured by expression data alone. Together, our benchmarking establishes a practical framework for clustering patient-specific GRNs and demonstrates the potential of GRN-derived representations for identifying clinically and biologically relevant tumor heterogeneity.

## Methods

### RNA-Seq Data Processing

RNA sequencing data for 33 cancer types (see Supplemental Table S1 for abbreviations and full names) were obtained from The Cancer Genome Atlas through the Genomic Data Commons using the TCGAbiolinks R package (version 2.25.3) (Colaprico et al. 2016). For each cancer type, gene-level raw count data were generated using the GDC “STAR – Counts” workflow and then downloaded. The STAR – Counts workflow is a pipeline used to quantify gene expression from RNA-seq data. It performs quantification using STAR alignments of tumor or normal samples and produces transcriptome profiling data (Dobin et al. 2013). The data were processed into SummarizedExperiment objects using the GDCprepare function, which integrates gene expression counts with associated sample-level and gene-level annotations. From these objects, raw count matrices (genes × samples) and corresponding metadata, including clinical identifiers, were extracted. Only primary tumor samples were retained for downstream analyses, resulting in a final processed dataset of 9,778 primary tumor samples.

Gene annotations provided by the GDC were used to filter the dataset to include only protein-coding genes when the gene type annotation was available, thereby reducing potential noise from non-coding transcripts. Raw counts were group-aware normalized using SNAIL to preserve co-expression structure (Hsieh et al. 2023). Batch effects were assessed using PCAplus (MBatch) (version 2.1.0), and corrected with sva (version 3.35.2) when necessary (Zhang et al. 2024). Detailed preprocessing and quality control procedures are described in Belova et al. (2025).

### Inference of Patient-Specific Gene Regulatory Networks

Patient-specific gene regulatory networks were generated on the pre-processed RNA-seq data following the methodology described in Belova et al. (2025). The SPONGE (Simple Prior Omics Network GEnerator) tool (Hovan and Kuijjer 2025) was used to construct prior regulatory and protein–protein interaction (PPI) networks by integrating transcription factor binding motifs from JASPAR (Rauluseviciute et al. 2024) and PPIs from STRING (Szklarczyk et al. 2023). The resulting prior networks were intersected with the gene expression dataset to retain only genes and transcription factors for which expression data were available. The filtered priors were then used as input to PANDA (Glass et al. 2013), which produced a complete bipartite reference graph based on the entire dataset, representing inferred transcription factor–gene regulatory relationships weighted by interaction likelihoods. The PANDA “aggregate” gene regulatory network was estimated based on a total of 9,778 primary tumor samples, for 19,034 genes, and 671 transcription factors. Individual, sample-specific regulatory networks were subsequently derived from this aggregate model by using LIONESS in combination with PANDA (Kuijjer et al. 2019).

### Input Data for Clustering

Clustering was applied to three data representations: gene expression data, indegree features, and edge-weight features. The gene expression and indegree representations each contained 19,034 features, corresponding to the number of genes retained in the final dataset. The edge-weight representation contained 12,771,814 features, corresponding to all possible transcription factor–gene interactions (671 transcription factors × 19,034 genes). The indegree, also called “Gene Targeting Score,” of a gene is defined as the sum of incoming regulatory edge weights from all transcription factors, reflecting the overall regulatory input received by that gene (Weighill et al. 2021).

### Consensus Clustering

Cola (version 2.14.0) systematically compares 20 combinations of four feature selection algorithms—coefficient of variation (CV), standard deviation (SD), median absolute deviation (MAD), and ability to correlate to other rows (ATC)—and five clustering algorithms—hierarchical clustering (hclust), model-based clustering (mclust), partitioning around medoids (PAM), k-means clustering (kmeans), and spherical k-means (skmeans). ATC is a feature selection approach introduced within the cola package. Detailed methodology and theoretical background of these feature selection and clustering algorithms can be found in the original cola publication (Gu et al. 2021).

#### Feature Selection Benchmarking

We first assessed the behavior of the four cola feature selection methods. This analysis was performed on four representative cancer types: ACC, GBM, ESCA, and KICH, to provide a representative subset, while keeping computational requirements manageable, as these datasets are not among the largest in TCGA, but provide sufficient sample sizes for initial profiling.

Runtime and memory usage were measured for each feature selection method in combination with each clustering algorithm, using both indegree and edge-weight data. This initial benchmarking was used to characterize the practical behavior of the candidate methods before scaling the analysis to all cancer types.

#### Consensus Clustering Parameters

Clustering analyses were performed using the consensus partition() function from the cola package with the default parameters. Consensus clustering was conducted by repeatedly subsampling a proportion of features (`p_sampling` = 0.8) and applying the partitioning methods across a range of cluster numbers (*k* = 2–6). For each configuration, 50 resampled partitions were generated (`partition_repeats` = 50) to construct a consensus matrix and derive stable cluster assignments.

For the largest dataset (BRCA), the `ATC_approx` function implemented in cola was used to approximate the ATC feature-selection step, applying 10% downsampling of samples during the consensus clustering procedure to reduce computational cost.

#### Consensus Clustering Performance Evaluation

Clustering performance was evaluated for all combinations of feature-selection methods and clustering algorithms across 33 TCGA cancer types using indegree and gene expression data. Edge-weight data were evaluated for the four representative cancer types described above, as time and memory profiling indicated that evaluating all 20 combinations across all cancer types would be computationally demanding and impractical at scale.

Performance was assessed using three metrics, each ranging from 0 to 1: mean silhouette score, (1*−*PAC), and concordance. An overall score was computed for each method combination as the average of the three metric values. The method combination achieving the highest overall score from this benchmarking analysis (with a score of 1 representing optimal performance) was selected for subsequent clustering analyses.

Cluster balance was additionally assessed using normalized entropy of the cluster-size distribution. For a clustering solution with k clusters, normalized entropy was calculated from the proportion of samples assigned to each cluster, with a value of 1 indicating equally sized clusters and lower values indicating increasingly imbalanced cluster assignments.

#### Selection of the Number of Clusters

The optimal number of clusters was determined using the `suggest best k()` function implemented in cola. This function applies a rule-based strategy to select the most appropriate value of *k*. First, values of *k* with a Jaccard index greater than 0.95 were excluded, because increasing *k* in these cases was considered not to provide sufficient additional information. If all candidate values of *k* were excluded under this criterion, no subgroup structure was considered detected.

Among the remaining candidate values, if any value of *k* had a 1 *−* PAC score greater than 0.9, the largest such *k* was selected as the optimal number of clusters, while other qualifying values were marked as optional. If this criterion was not met, the optimal *k* was determined by majority vote among the value of *k* with the highest 1 *−* PAC score, highest mean silhouette width, and highest concordance (Gu et al. 2021).

#### Proportion of Features Retained by SD-Based Cola Selection

After selecting SD + k-means as the final clustering strategy, we examined the proportion of features retained by the SD feature selection method. To determine the optimal number of features to retain, cola applies an elbow-based approach to the ranked SD values (Gu and Hübschmann 2022), with an upper bound of 10% of the total features. Specifically, features are ordered by decreasing SD and the elbow point of the SD distribution curve is identified. This point represents where the inclusion of additional features yields diminishing increases in variability. Features above this elbow threshold are retained to construct the submatrix, after which row-wise z-score scaling is applied prior to downstream clustering.

#### Comparison of Edge-Weight and Indegree Clustering

To assess the consistency of clustering results derived from different network representations, we compared cluster assignments obtained from edge-weight matrices and indegree matrices using the Adjusted Rand Index (ARI). The ARI measures the agreement between two clustering solutions while adjusting for similarity that could occur by chance. The index ranges from -1 to 1, where a value of 1 indicates perfect agreement between clusterings, 0 corresponds to random agreement, and negative values indicate less agreement than expected by chance. Higher ARI values therefore indicate greater similarity between clustering structures derived from the two representations.

### Clinical Variables Association with Clusters

Clinical metadata were retrieved from TCGAbiolinks (version 2.36.0) for samples from The Cancer Genome Atlas. Cluster assignments obtained from the consensus clustering analysis were matched to patient clinical records using the TCGA submitter barcode (first 12 characters), allowing cluster membership to be integrated with available demographic and clinical information. In total, 211 clinical variables were collected across all cancer types, although not every variable was present in each cancer dataset. Administrative identifiers, technical metadata, and treatment-related variables not relevant to baseline clinical characterization were excluded prior to analysis. Clinical variables were tested only when sufficient data were available within each cancer type: variables were required to have non-missing values in at least two clusters, with a minimum of three samples per cluster. In addition, categorical variables were required to have at least two observed categories with a minimum of three samples per category, whereas numerical variables were required to have at least two unique observed values. Variables that did not meet these criteria were excluded from the corresponding cancer-specific analysis. All statistical analyses were performed using only samples with non-missing values for the variable under investigation.

Associations between cluster membership and categorical clinical variables were assessed using chi-squared tests. For numerical variables, one-way ANOVA was used when the data followed a normal distribution according to the Shapiro–Wilk test, otherwise, Kruskal–Wallis tests were applied. P-values were adjusted for multiple testing using the Benjamini–Hochberg false discovery rate (FDR) procedure (Benjamini and Hochberg 1995), separately to categorical and numerical results, and significant variables were identified at FDR *<* 0.05.

### Pathway Enrichment Analysis

To assess the biological processes associated with clusters obtained from the three feature representations (indegree, edge-weight, and gene expression), we performed pathway enrichment analysis using both the indegree and gene-expression datasets. Because edge-weight matrices do not provide per-gene measurements, they cannot be used directly for GSEA. Instead, we used the sample-level cluster assignments derived from the three feature representations and evaluated them only in the two datasets containing per-gene quantities (expression and indegree). Thus, for any clustering modality—including edge-weight-based clustering—samples were grouped according to their assigned cluster, and differential analyses were performed on gene expression or indegree values for those same samples. Pathway enrichment was therefore always computed on per-gene measurements, independent of the feature space used to generate the clusters.

Within each evaluation dataset (indegree or expression), we then performed differential analysis using limma (version 3.64.3), constructing a one-vs-rest contrast for every cluster (Ritchie et al. 2015). The resulting linear models yielded moderated t-statistics for all genes, summarizing how strongly each gene distinguished a given cluster from the rest in that dataset. These t-statistics were ranked to generate ordered gene lists, which served as input for Gene Set Enrichment Analysis (GSEA) (Subramanian et al. 2005; Mootha et al. 2003). Gene set enrichment was performed using Reactome pathway annotations (C2:CP:REACTOME) obtained via the msigdbr R package (version 25.1.1) for Homo sapiens (Dolgalev 2025). Pathway enrichment was performed using 10,000 permutations, and pathways were considered significant if they satisfied within-contrast Benjamini–Hochberg correction (*p*_adj_ *<* 0.05), exhibited an enrichment score with magnitude greater than 0.5 (|*ES*| *>* 0.5), and a minimum size of 10 genes (Fig. S3).

### Survival Associations of Clustering Results

Finally, survival differences between clusters were assessed using Kaplan–Meier analysis implemented through the `TCGAanalyze_survival` function from the `TCGAbiolinks` R package. For each cancer type, tumor samples were grouped according to their cluster assignments obtained from consensus clustering of the three data modalities (indegrees, edge weights, and gene expression). Clinical data were retrieved from The Cancer Genome Atlas (TCGA) using `GDCquery_clinic`, in the same manner as in the clinical variable association analysis.

Within the clinical dataset, the `vital_status` column indicates whether a patient was alive or deceased at the time of the last follow-up. Survival time was derived from TCGA clinical variables, using `days_to_death` for deceased patients, representing the number of days from the initial diagnosis to the patient’s death, and `days_to_last_follow_up` for patients who were still alive, representing the time between the initial diagnosis and the most recent clinical follow-up visit. Kaplan–Meier survival curves were generated for each cluster, and statistical differences between groups were evaluated using the log-rank test. Associations were considered statistically significant at *p <* 0.05.

### Computational Environment

All computational analyses were performed on the CSC Finnish national high-performance computing infrastructure using the Puhti and Mahti systems. Clustering analyses involving large matrices were primarily executed on Puhti using the high-memory `hugemem` partition, with 1496 GB RAM and 40 CPU cores, to accommodate the memory requirements of edge-weight features. Additional analyses with gene-expression and indegree-based clustering were performed on Mahti using the `medium` partition with 200 GB of allocated memory and four CPU cores per job. All jobs were submitted through the SLURM workload manager and executed in R version 4.5.0 using cola version 2.14.0.

The clustering benchmark and associated downstream analyses were implemented in BOAR (Benchmarking Of Approaches for Regulatory-Network Clustering), a reproducible R analysis pipeline developed for this study.

## Results

### Benchmarking Feature Selection and Clustering Methods for GRN-Derived Features

To identify a robust strategy for clustering patient-specific GRNs, we first evaluated eleven clustering packages in R and Python. Because our objective was to cluster patients across a cohort, rather than to identify communities of genes or nodes within a single network, we focused the final comparison on methods that can operate on numeric sample-level feature matrices (see Supplemental Table S2). Our evaluation criteria included the ability to handle high-dimensional data, support for feature selection, provision of diagnostics for cluster quality assessment, and computational efficiency. Based on this comparison, we identified the cola package (Gu et al. 2021) as the most suitable tool for clustering GRNs. This framework performs consensus clustering by repeatedly clustering randomly sampled subsets of the data, thereby summarizing clustering robustness and producing a consensus classification of all samples.

### SD + K-Means Provides Strong Clustering Performance and Computational Feasibility

We evaluated all feature selection methods and clustering algorithms available in the cola framework, comprising four feature selection methods (SD, MAD, CV, ATC) and five clustering algorithms (kmeans, skmeans, hclust, pam, and mclust). Expression and indegree data were assessed across all 33 TCGA cancer types, whereas edge-weight data were evaluated in four representative cancer types because of the substantially higher dimensionality and computational burden of this representation.

Clustering performance was compared using three metrics: mean silhouette, (1 *−* PAC), and concordance. Mean silhouette evaluates how well samples fit within their assigned clusters relative to neighboring clusters. The (1 *−* PAC) score reflects consensus clustering stability, with higher values indicating fewer ambiguously clustered sample pairs. Concordance measures the agreement between individual clustering partitions and the final consensus solution. To summarize these metrics, we calculated an overall performance score for each method combination as the average of mean silhouette, (1 *−* PAC), and concordance.

Across the GRN-derived features, the combination of standard deviation (SD) for feature selection and k-means for sample clustering consistently achieved the strongest overall performance (Fig. 2). SD ranks features according to their variability across samples, thereby retaining features with high variance that may capture biologically meaningful differences between samples. K-means then partitions samples into *k* clusters by iteratively assigning each sample to the nearest cluster centroid and minimizing the within-cluster sum of squared distances. For edge-weight features specifically, SD + k-means ranked first among all tested combinations, achieving the highest overall score (0.997). For indegree features, SD + k-means again showed the best overall ranking, outperforming the other method combinations with an overall score of 0.985. Expression-based clustering also showed competitive performance, although its top-ranked combination was ATC + k-means (0.976) rather than SD + k-means (Fig. 2A).

**Figure 1.**
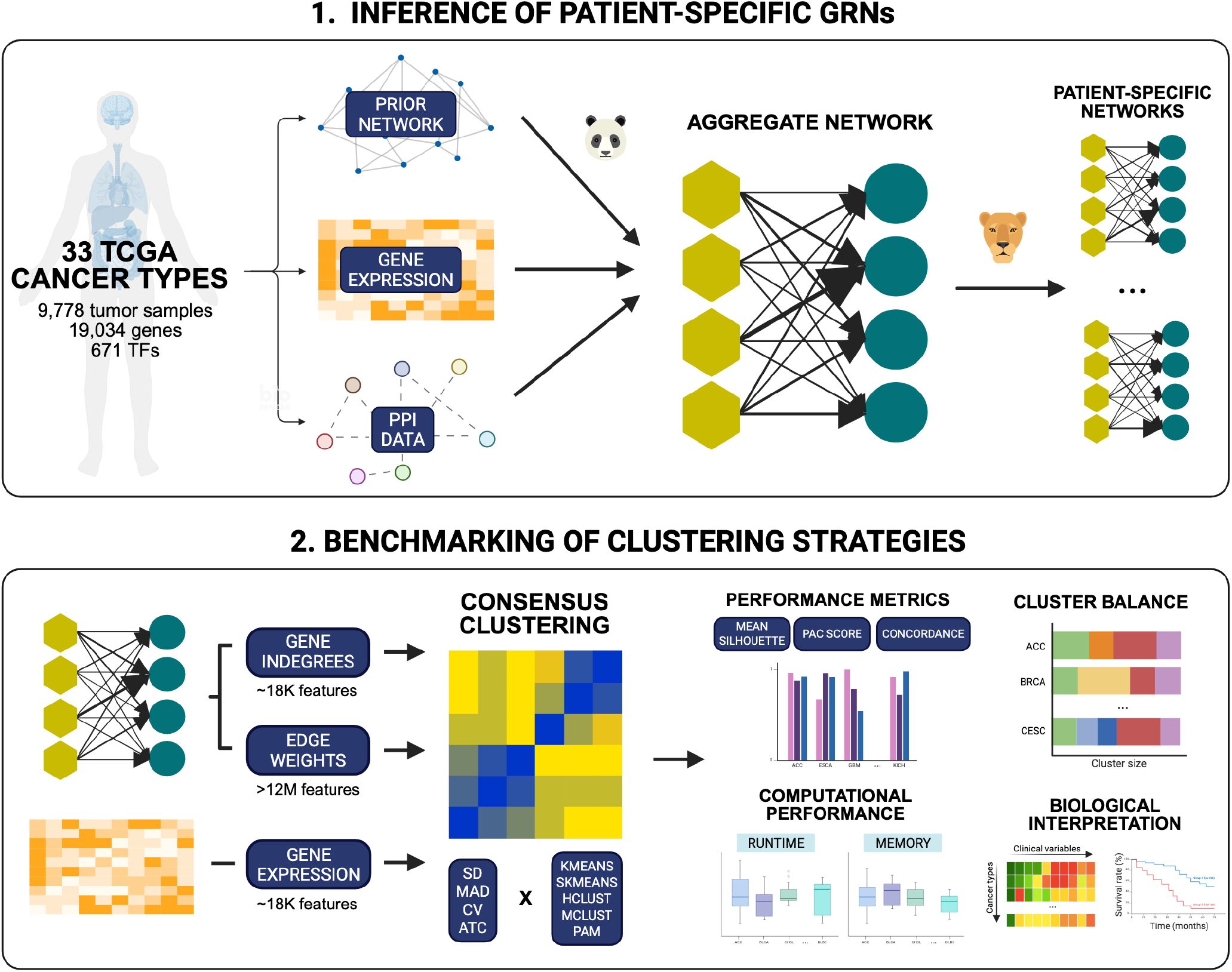
Overview of the approach; 33 TCGA cancer types were used to infer patient-specific GRNs and three different representations were used to benchmark 20 clustering strategies.

**Figure 2.**
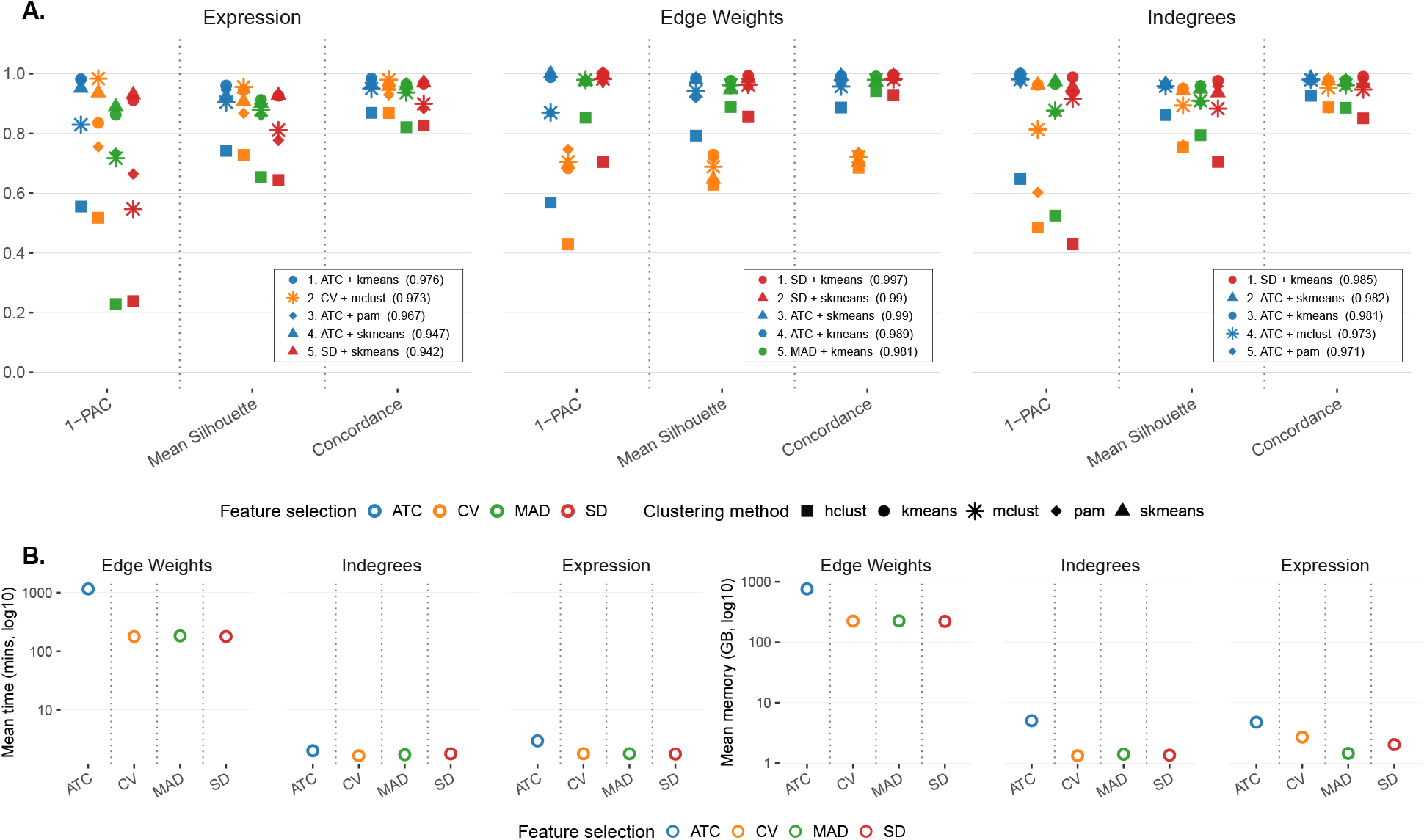
Benchmarking clustering performance and computational feasibility across feature-selection and clustering-method combinations. (A) Clustering performance for gene expression, edge-weight, and indegree features using three evaluation metrics: 1–PAC, mean silhouette, and concordance. Expression and indegree features were evaluated across all 33 TCGA cancer types, whereas edge-weight features were evaluated in four representative cancer types given the substantially larger number of features and associated computational requirements. Points represent combinations of feature-selection methods and clustering algorithms implemented in the cola framework, with colors indicating feature-selection methods and shapes indicating clustering algorithms. Insets list the five highest-ranked method combinations for each data representation. (B) Computation time (left) and mean memory usage (right) on a log_10_ scale across feature-selection methods.

Because edge-weight representations contain substantially larger numbers of features than gene expression and indegree data, we also evaluated computation time and memory usage for all 20 method combinations across four representative cancer types and the three data modalities (Fig. 2B). The ATC feature-selection method was consistently the most computationally demanding, particularly for edge-weight data. On a logarithmic scale, ATC required approximately an order of magnitude more time and memory than the other methods. This difference is especially relevant for edge-weight datasets, which contain 12,771,814 features, compared with 19,034 features in both the indegree and gene expression representations. These results highlight the substantial computational burden associated with clustering full edge-weight representations and demonstrate that the choice of feature-selection strategy strongly influences computational scalability.

Taken together, these benchmarking results identified SD feature selection combined with k-means clustering as the best-performing strategy for clustering patient-specific GRNs. We therefore used this strategy for all subsequent analyses.

### SD + K-Means Produces Stable GRN-Derived Clusters Across Cancer Types

Using the selected SD + k-means strategy, we next evaluated clustering performance across cancer types for the GRN-derived representations. Edge-weight and indegree features both produced consistently high clustering-quality metrics, with 1– PAC, concordance, and mean silhouette values concentrated near 1 across most cancer types. The distributions were broadly similar between the two GRN modalities, although indegree-based clustering appeared slightly more stable, with fewer outliers across the three metrics. By comparison, gene expression-based clustering showed a wider spread in clustering-quality scores (Fig. 3A).

**Figure 3.**
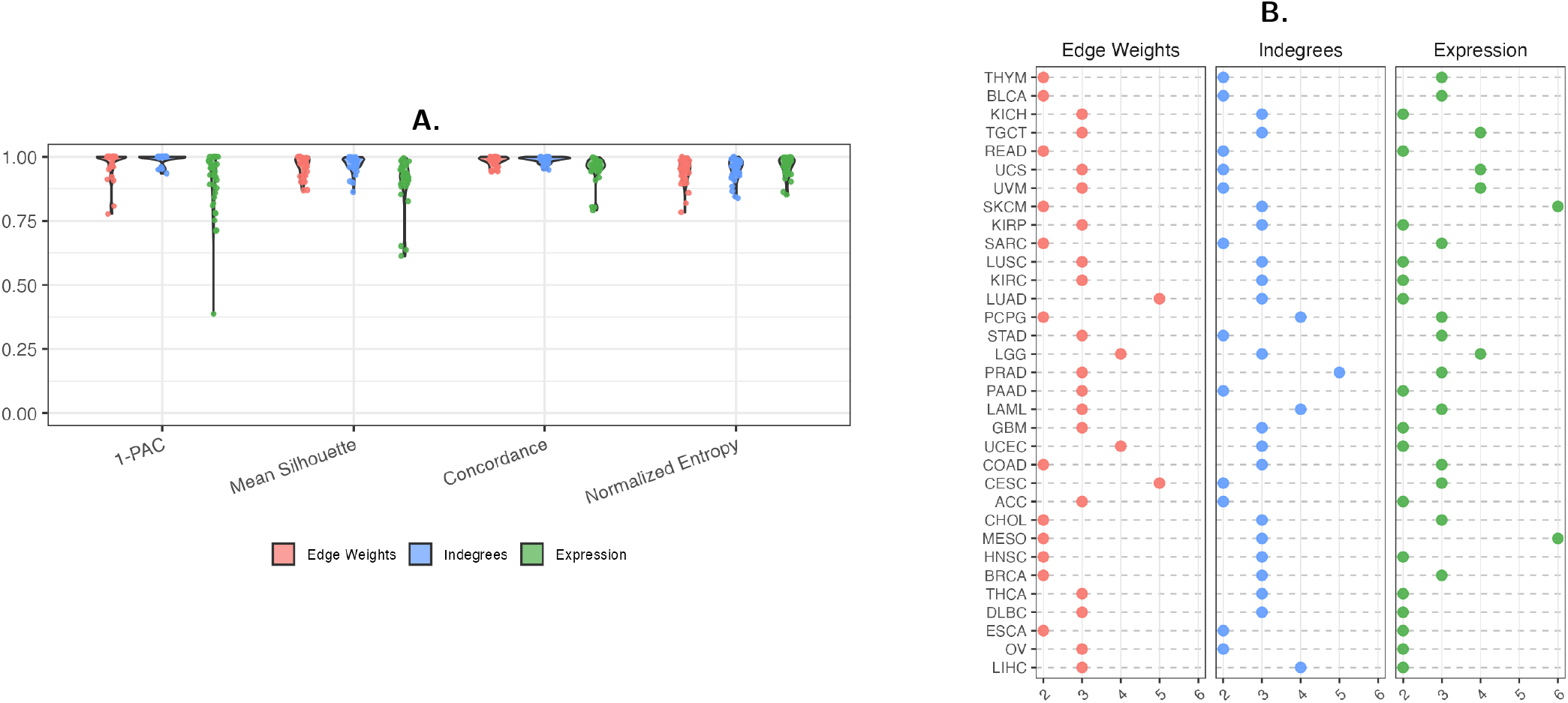
(A) Violin plots showing the distributions of 1–PAC, mean silhouette, concordance, and normalized entropy scores across cancer types. Individual points represent cancer-specific clustering results for edge-weight, indegree, and expression representations. (B) Selected number of clusters for each cancer type using the rule-based `suggest_best_k()` procedure across edge-weight, indegree, and expression modalities.

We then assessed cluster balance for each cancer type and data modality using normalized entropy, where a value of 1 indicates equally sized clusters. Across gene expression, edge-weight, and indegree representations, normalized entropy values were generally high (Fig. 3A). Gene expression-based clustering showed the most balanced cluster assignments overall, with values concentrated close to 1 and an average normalized entropy of 0.962. GRN-derived representations also showed high cluster balance, with edge-weight and indegree features achieving average normalized entropy values of approximately 0.942 and 0.949, respectively. Edge-weight clusters showed slightly greater variability and included several lower-balance cases, whereas indegree clusters were more tightly distributed near the upper range. The sample distribution across clusters for each cancer type and data modality is shown in Supplemental Figure S1.

Finally, we examined the number of subgroups selected for each cancer type using the rule-based `suggest_best_k()` procedure (Fig. 3B). This procedure excludes candidate values of *k* that do not provide substantially new clustering structure, then selects the largest remaining *k* with strong stability based on (1 *−* PAC). The selected number of clusters was generally small across both GRN-derived modalities, with most cancer types assigned either two or three subgroups. Across the 66 total GRN-derived representations evaluated, 25 selected *k* = 2 and 33 selected *k* = 3, whereas only 5 selected *k* = 4 and 3 selected *k* = 5; no representation selected *k* = 6. Although the edge-weight and indegree modalities often selected similar values of *k*, the selected number of subgroups differed between modalities for 21 of the 33 cancer types, indicating that the two GRN-derived feature spaces capture partially distinct clustering structure.

### Indegree and Edge-Weight Clusters Show Partial Concordance Across Cancer Types

To assess agreement between clustering partitions derived from the two network representations, we computed the adjusted Rand index (ARI). ARI measures the similarity between two clustering partitions while correcting for agreement expected by chance, with values ranging from 0, indicating random agreement, to 1, indicating perfect alignment. Although ARI values varied substantially across cancer types, all values were greater than zero, indicating higher-than-random similarity between edge-weight- and indegree-based clustering solutions.

The three cancer types with the highest agreement between edge-weight and indegree clustering assignments were thymoma (THYM; ARI = 0.838), bladder urothelial carcinoma (BLCA; ARI = 0.765), and kidney chromophobe (KICH; ARI = 0.726), each with ARI values greater than 0.7. For these three cancer types, the number of detected subgroups was identical across the two modalities, with two subgroups identified for THYM and BLCA and three subgroups identified for KICH. This concordance does not appear to be driven by cohort size, as KICH is among the smaller cohorts, whereas THYM and BLCA represent moderate-sized cohorts.

Across all cancer types, the mean ARI between edge-weight- and indegree-based clustering solutions was 0.355. In contrast, comparisons between GRN-derived and gene expression-based clustering produced substantially lower mean ARI values, with a mean ARI of 0.096 for edge weights versus expression and 0.066 for indegrees versus expression (Fig. 4). These results suggest that indegree representations retain part of the information present in full edge-weight networks, while both GRN-derived representations capture clustering patterns that are distinct from those obtained using gene expression alone.

**Figure 4.**
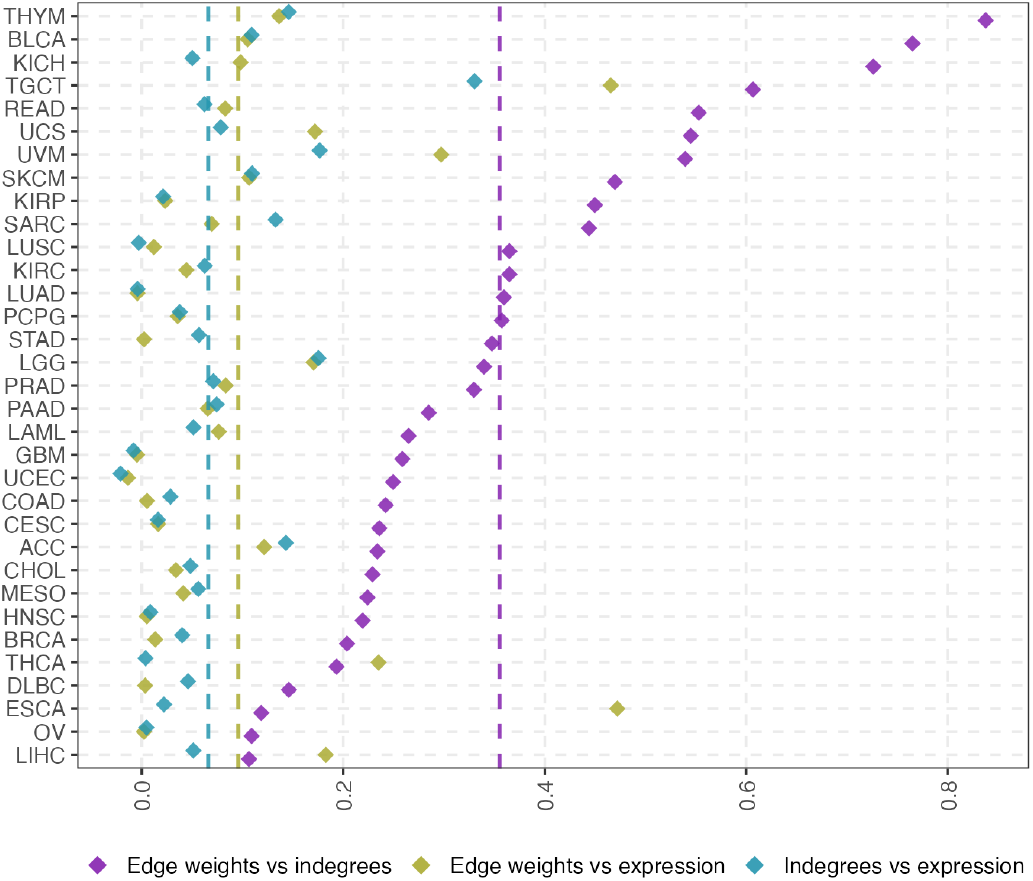
Concordance between clustering assignments derived from edge-weight, indegree, and gene expression representations across cancer types. Each point represents the adjusted Rand index (ARI) between two clustering solutions for a given cancer type using the SD + k-means strategy. Three pairwise comparisons were performed: edge weights and indegrees, edge weights and gene expression, and indegrees and gene expression, yielding mean ARI values of 0.355, 0.096, and 0.066, respectively. Cancer types are ordered by ARI values for the edge-weight–indegree comparison. Dashed vertical lines indicate the mean ARI for each comparison.

### Network-Derived Clusters Capture Complementary Clinical Associations

Having observed both shared and distinct clustering patterns across data representations, we next tested associations between cluster membership and available clinical variables, considering both numerical and categorical variables (see Methods), since molecular subgroups may reflect clinically relevant patient heterogeneity. Such associations should be interpreted as supporting evidence of clinical relevance rather than as a gold standard, as unsupervised clustering may reveal molecular heterogeneity that is not captured by existing clinical annotations.

Across cancer types, a total of 21 clinical variable–cancer type combinations were significantly associated with clusters derived from both GRN-derived representations, whereas 20 were uniquely associated with indegree-based clusters and 25 were uniquely associated with edge-weight-based clusters (Fig. 5). This indicates that the two GRN-derived representations capture both common and complementary aspects of clinically relevant tumor heterogeneity. The observed overlap was greater than expected by chance (*p <* 2.2 *×* 10^*−*16^; odds ratio = 37.7; 95% CI: 16.8–86.5).

**Figure 5.**
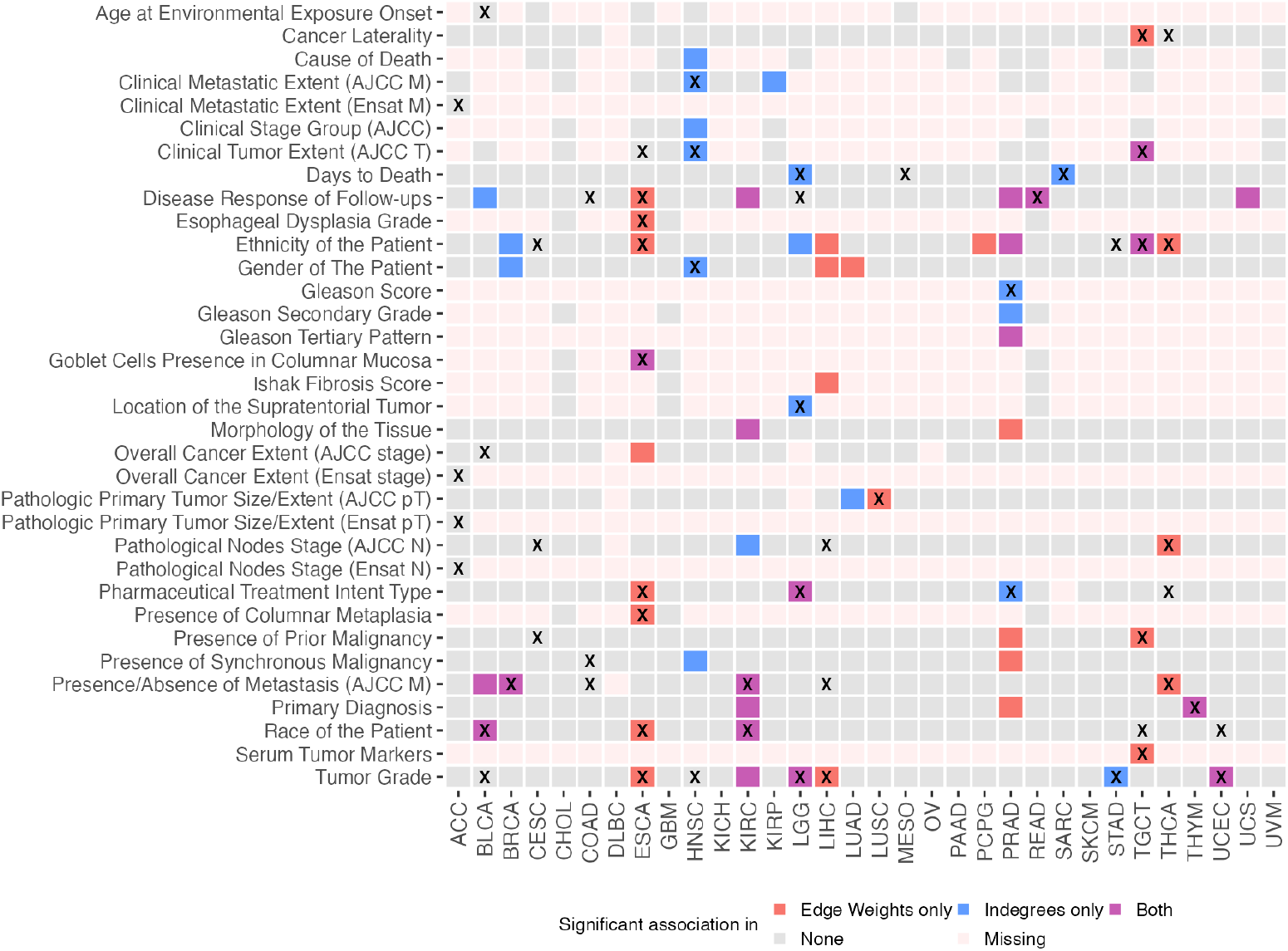
Clinical associations of GRN-derived clustering assignments across TCGA cancer types. Rows represent clinical variables and columns represent cancer types. Cell colors indicate whether subtype assignments were significantly associated with the corresponding clinical variable for edge-weight-based clusters only, indegree-based clusters only, both GRN-derived representations, or neither representation. Pale pink cells indicate clinical variables that were missing or unavailable for a given cancer type. An “X” marks clinical variables that were significantly associated with gene expression-based clustering assignments.

We also compared the clinical associations detected using GRN-derived clusters with those detected using gene expression-based clusters. Overall, 30 clinical variable–cancer type combinations were identified by at least one network-derived representation but not by gene expression, whereas 24 were detected by gene expression alone. In addition, 36 associations were shared between gene expression and at least one network-derived representation.

Several of the network-specific associations involved clinically meaningful variables. In particular, network-derived clusters identified associations with six clinical variables that were not detected in any cancer type using expression-based clusters: cause of death, clinical stage group, Gleason secondary grade, Gleason tertiary pattern, Ishak fibrosis score, and tissue morphology. Among these, tissue morphology was especially notable because it was associated with network-derived clustering in two cancer types, kidney renal clear cell carcinoma (KIRC) and prostate adenocarcinoma (PRAD), whereas each of the other five variables was associated with network-derived clustering in only one cancer type. KIRC and PRAD were particularly enriched for clinical associations identified by network-derived clustering. In KIRC, seven clinical associations were detected, six of which were shared by both network representations. Four of these were not detected using expression-based clustering and include, for example, tumor grade. In PRAD, ten clinical associations were detected. Only two of these were also identified in expression-based clustering. The added value of GRN-derived clustering was also apparent within individual cancer types. Four cancer types, KIRP, lung adenocarcinoma (LUAD), pheochromocytoma and paraganglioma (PCPG), and uterine carcinosarcoma (UCS), showed clinical associations only for network-derived clusters and not for expression-based clusters. Among these, LUAD had two network-specific clinical associations, whereas the other three each had one.

### Network-Derived Clusters Show Prognostic Associations in Selected Cancer Types

Recent studies have demonstrated that gene indegrees can be informative for identifying survival-associated patterns (Pop et al. 2025). Thus, we hypothesized that downstream survival analysis of the identified clusters can provide insight into prognostic subgroup structure and highlight biological or regulatory processes that contribute to cancer aggressiveness. To this end, we investigated whether these cluster-level differences were associated with patient survival. For each cancer type and data modality, overall survival was compared independently between the identified clusters using Kaplan–Meier survival analysis and log-rank tests. Figure 6 summarizes the corresponding p-values for each cancer type and data representation.

**Figure 6.**
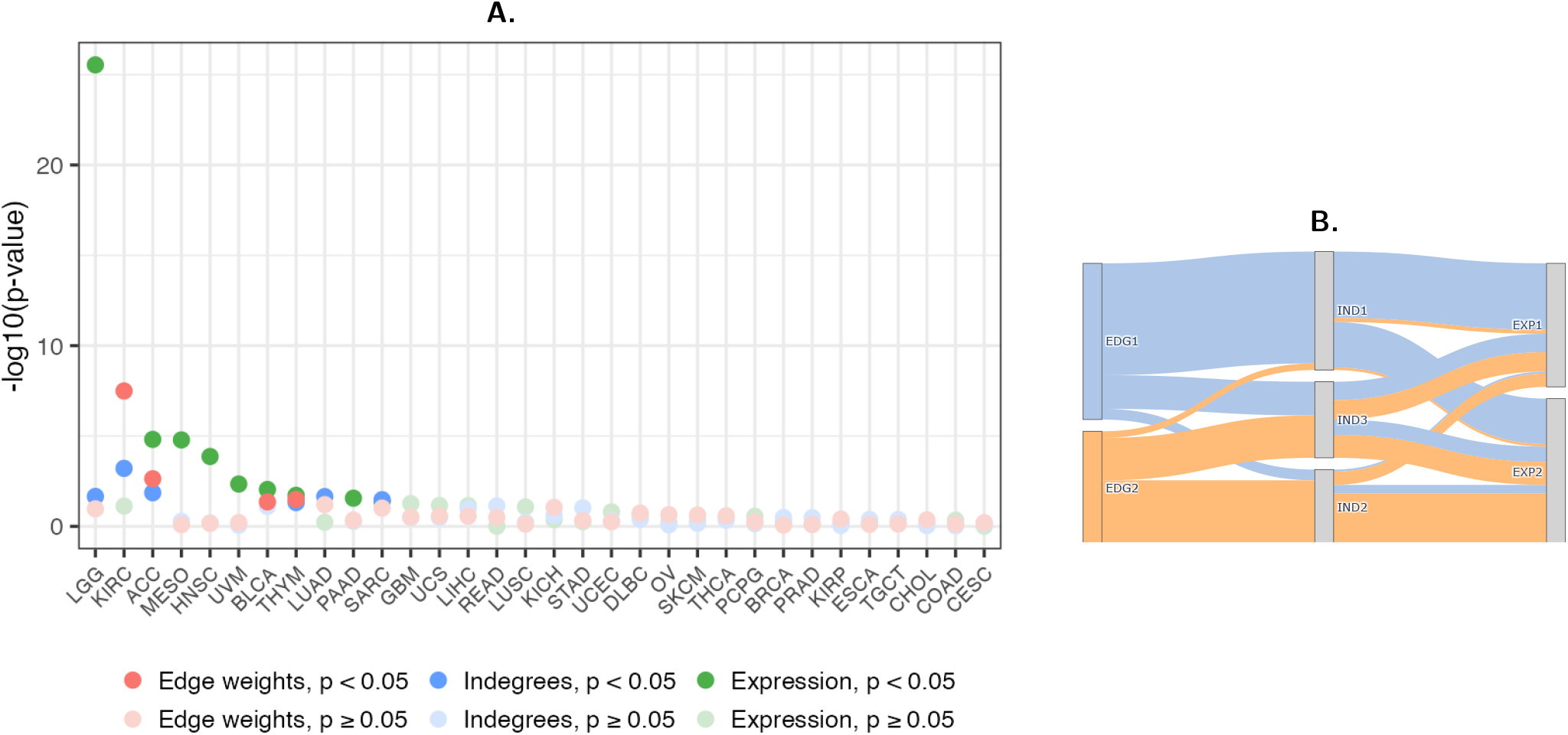
Survival associations and cluster correspondence across data modalities. (A) Survival analysis of clusters across cancer types and data modalities. Cancer types are ranked by *−* log_10_(*p*-values) in descending order. Red squares indicate gene expression, triangles indicate indegrees, and circles indicate edge weights; filled symbols denote significant associations with survival (*p <* 0.05). (B) Sankey plot illustrating cluster correspondence between edge-weight, indegree and gene expression clustering assignments for KIRC. Cluster sizes for gene expression were *n*_1_ = 243 and *n*_2_ = 290; for indegrees, *n*_1_ = 232, *n*_2_ = 149, and *n*_3_ = 149; and for edge weights, *n*_1_ = 306 and *n*_2_ = 224.

Using a significance threshold of *p <* 0.05 for each cancer-specific analysis, survival differences were observed in eleven cancer types across the three data modalities. Network-derived clusters showed significant survival differences in seven cancer types, whereas gene expression–derived clusters showed significant differences in eight. Four cancer types showed significant survival differences only for gene expression–derived clusters, whereas two showed significant differences only for network-derived clusters. Among the seven cancer types with significant network-based associations, the corresponding log-rank p-values were lower than those obtained for gene expression–derived clusters in three cancer types: KIRC, LUAD, and sarcoma (SARC).

Notably, KIRC was the only cancer type that showed significant survival associations for both edge-weight- and indegree-derived clusters, while gene expression-derived clusters did not reach significance. For edge-weight, indegree, and gene expression clusters, respectively, the log-rank p-values were *<* 0.0001, 0.00062, and 0.076 (Supplemental Figure S2). Comparison of the cluster assignments indicated partial overlap among the patient groups identified by the three data modalities (Figure 6B), suggesting that the edge-weight-derived clustering captured different patterns from those based on gene expression or indegree profiles.

To illustrate how biologically meaningful the network-based stratification could be, we focused on KIRC, where clustering based on edge weights showed the strongest association with survival. We then asked whether these clusters also reflected differences in pathway activity. To address this, we performed GSEA using Reactome pathway annotations. Because edge weights are defined between transcription factors and genes rather than as per-gene measurements, they cannot be used directly to rank genes for GSEA. We therefore retained the KIRC edge-weight cluster labels to define the patient groups, but ranked genes using the corresponding indegree and gene expression profiles. For each subgroup, genes were ranked according to their moderated t-statistics from the one-vs-rest differential analysis of indegree or expression, and the resulting ranked lists were used as input for GSEA (Supplemental Figure S3).

Across these KIRC subgroups, GSEA identified 128 unique enriched Reactome pathways using indegree-based gene rankings and 98 using gene-expression-based rankings (Supplemental Figure S4). We then focused on the ten pathways with the lowest enrichment p-values to summarize the strongest biological signals (Supplemental Table S3). These pathways showed an opposing biological pattern between the two edge-weight-derived KIRC clusters. Cluster 1, which was associated with shorter survival, was enriched for pathways related to cell-cycle progression, DNA replication, mitotic control, and renal homeostasis. Dysregulation of cell-cycle and DNA replication programs is consistent with increased proliferative capacity and may reflect more aggressive tumor behavior. In contrast, cluster 2, which was associated with longer survival, showed enrichment for immune-related pathways, suggesting that stronger immune activity may characterize patients with more favorable outcomes. Together, these results indicate that network-derived clustering can identify clinically relevant patient subgroups and highlight biological programs associated with differences in survival.

## Discussion

In this study, we systematically benchmarked feature-selection and clustering strategies for patient-specific GRNs across 33 TCGA cancer types. We found that SD feature selection combined with k-means clustering provided the best balance between clustering quality and computational scalability for GRN-derived representations. In contrast, ATC-based feature selection was less informative for network-derived features, despite previously reported improvements for gene expression datasets (Gu et al. 2021), and required substantially greater time and memory, particularly for edge-weight data. This may be because SD directly prioritizes patient-to-patient variability, whereas ATC depends on feature–feature correlation structure, which may behave differently in high-dimensional GRN-derived matrices. The strong performance of k-means may partly result from its compatibility with SD-based feature selection, because selecting features with high variability across patients emphasizes the differences used by k-means to separate samples into compact, centroid-based clusters.

We also examined differences in clustering results across gene expression, indegree, and edge-weight representations, which produced somewhat overlapping but non-identical subgroup structures. Indegree- and edge-weight-based clustering achieved comparable cluster quality across cancer types, despite the much higher dimensionality of the edge-weight matrices. Their subgroup assignments also showed partial overlap, indicating that indegrees retain some of the clustering-relevant information present in the full network representation. At the same time, gene expression- and network-derived clusters showed distinct patterns of clinical, pathway, and survival associations across cancer types, suggesting that the different representations capture complementary aspects of tumor heterogeneity. The greater detail provided by edge-weight representations, however, comes at a substantial computational cost in terms of both time and memory. Indegrees therefore offer a more compact network summary that remains informative while being considerably more scalable. When computational resources are limited, indegree-based clustering may thus provide a practical alternative to clustering based on the full edge-weight representation.

Because unsupervised clustering lacks a definitive ground truth in real cancer cohorts, we evaluated clustering strategies using complementary criteria, including internal clustering metrics, computational feasibility, and downstream clinical and biological interpretability. Clinical annotations, survival associations, and pathway enrichment should therefore be interpreted as supporting evidence that the clusters capture relevant tumor heterogeneity, rather than as definitive validation of a single correct clustering solution. In this context, clusters derived from gene expression, indegrees, and edge weights were associated with clinical variables and survival in selected cancer types. Some associations were detected only in network-derived clusters, suggesting that patient-specific GRNs capture aspects of tumor heterogeneity not fully represented by gene expression alone. KIRC provided an illustrative example in which edge-weight-derived clusters were associated with survival and showed opposing pathway-enrichment patterns, with cell-cycle and DNA-replication pathways enriched in the shorter-survival cluster and immune-related pathways enriched in the longer-survival cluster.

To our knowledge, no previous study has combined patient-specific gene regulatory network (GRN)-derived clustering with cola to perform subtype analysis separately within multiple cancer types. Previous studies have shown that incorporating gene regulatory network information can improve cancer subtyping compared with approaches based solely on gene expression (Xu et al. 2016; Nakazawa et al. 2021). However, these approaches relied on a single regulatory network shared across patients to guide subtype identification. In contrast, our framework uses PANDA and LIONESS to infer an individualized GRN for each patient, providing a sample-specific weighted regulatory architecture that captures inter-patient differences in regulatory organization within each cancer type.

Of course, our study has several limitations. The generation of patient-specific GRNs and the analysis of their edge weights are computationally demanding, which may reduce the feasibility of full network-based clustering for research groups with limited computational resources. In addition, our benchmarking was performed using single-sample GRNs inferred with PANDA and LIONESS. Although this provided a consistent framework for comparing clustering strategies across cancer types, the extent to which the conclusions generalize to other single-sample network inference methods or to networks derived from other data modalities remains to be established.

To support reproducibility and facilitate extensions of this work, we developed BOAR (Benchmarking Of Approaches for Regulatory-Network Clustering), a reusable R analysis pipeline that implements the clustering benchmark and associated downstream clinical and survival analyses described in this study. BOAR can be applied to independent datasets to reproduce and adapt the same analysis workflow. The same framework could also be used to benchmark clustering strategies for other types of single-sample network representations and in other biological contexts. For example, methods such as SCORPION enable the reconstruction of comparable GRNs from single-cell or single-nucleus RNA-seq data for population-level analyses and could support patient stratification based on regulatory networks modeled within individual cell types (Osorio et al. 2024). The framework may also be applicable beyond gene regulatory networks. For instance, microbial co-abundance networks have been used to characterize relationships among microbial taxa and analogous network-based representations could potentially support microbiome-based subtyping (Minot and Willis 2019). It may also support studies of heterogeneity in other complex diseases and biological systems in which inter-individual differences in regulatory or interaction patterns are important, including autoimmune disorders. In the present study, clinical variables were used after clustering to assess the clinical relevance of the identified subgroups. Future extensions could instead incorporate selected clinical variables directly into network inference or integrative clustering, allowing molecular and clinical information to jointly inform patient stratification.

## Acknowledgments

The authors wish to acknowledge CSC – IT Center for Science, Finland, for computational resources.

## Data Availability

The data analyzed in this study are publicly available through the National Cancer Institute Genomic Data Commons (GDC; https://portal.gdc.cancer.gov/), including data from The Cancer Genome Atlas (TCGA) and the Clinical Proteomic Tumor Analysis Consortium (CPTAC).

## Code Availability

BOAR (Benchmarking Of Approaches for Regulatory-Network Clustering), the reproducible R analysis pipeline supporting the clustering benchmark and associated downstream analyses presented in this study, is publicly available at https://github.com/kuijjerlab/BOAR. The repository includes the analysis code, a reproducible software environment, example input data, and documentation for running the workflow.

## Supplemental Tables and Figures

**Table S1.**
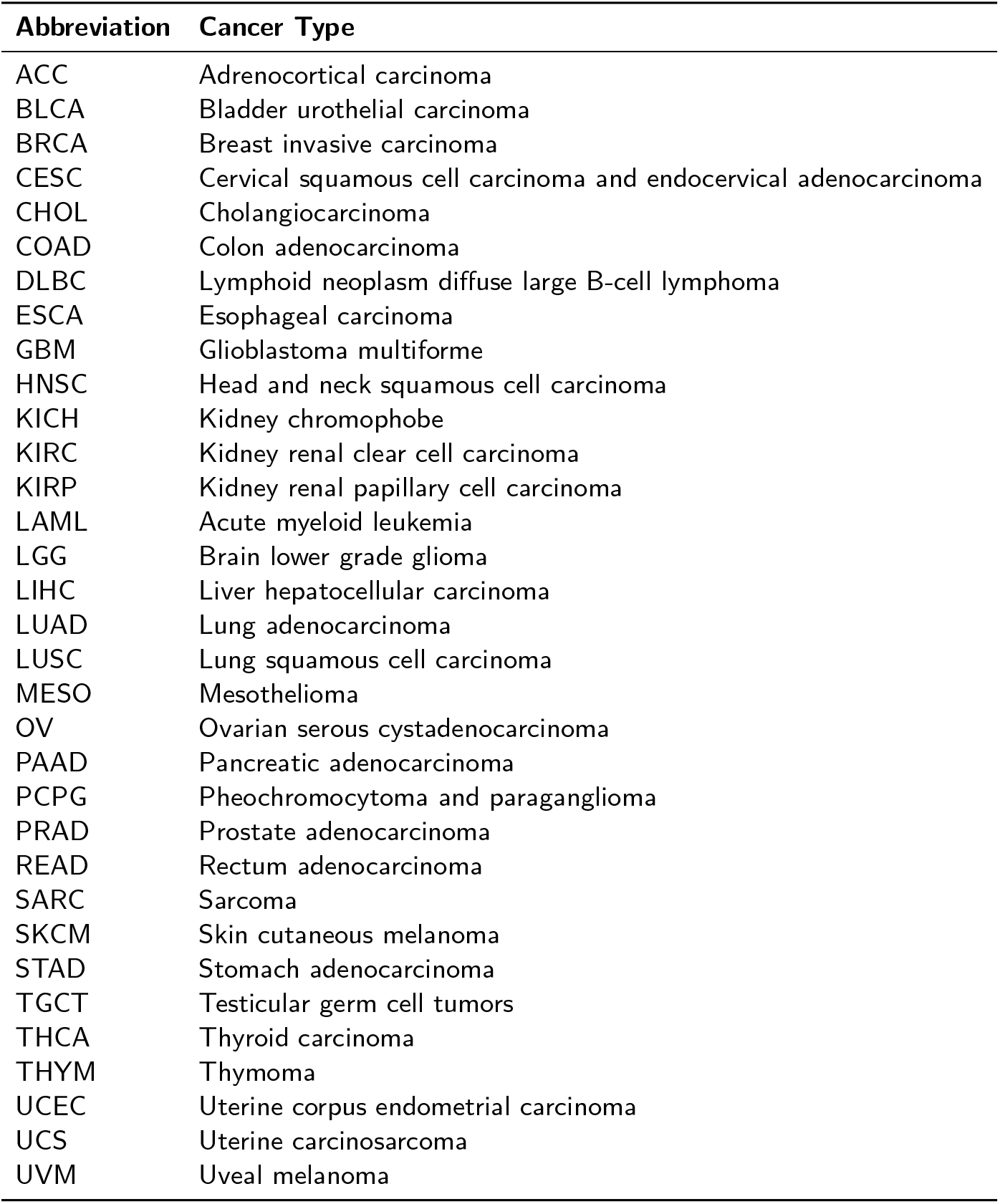
Cancer type abbreviations and their meanings.

| Abbreviation | Cancer Type |
| --- | --- |
| ACC | Adrenocortical carcinoma |
| BLCA | Bladder urothelial carcinoma |
| BRCA | Breast invasive carcinoma |
| CESC | Cervical squamous cell carcinoma and endocervical adenocarcinoma |
| CHOL | Cholangiocarcinoma |
| COAD | Colon adenocarcinoma |
| DLBC | Lymphoid neoplasm diffuse large B-cell lymphoma |
| ESCA | Esophageal carcinoma |
| GBM | Glioblastoma multiforme |
| HNSC | Head and neck squamous cell carcinoma |
| KICH | Kidney chromophobe |
| KIRC | Kidney renal clear cell carcinoma |
| KIRP | Kidney renal papillary cell carcinoma |
| LAML | Acute myeloid leukemia |
| LGG | Brain lower grade glioma |
| LIHC | Liver hepatocellular carcinoma |
| LUAD | Lung adenocarcinoma |
| LUSC | Lung squamous cell carcinoma |
| MESO | Mesothelioma |
| OV | Ovarian serous cystadenocarcinoma |
| PAAD | Pancreatic adenocarcinoma |
| PCPG | Pheochromocytoma and paraganglioma |
| PRAD | Prostate adenocarcinoma |
| READ | Rectum adenocarcinoma |
| SARC | Sarcoma |
| SKCM | Skin cutaneous melanoma |
| STAD | Stomach adenocarcinoma |
| TGCT | Testicular germ cell tumors |
| THCA | Thyroid carcinoma |
| THYM | Thymoma |
| UCEC | Uterine corpus endometrial carcinoma |
| UCS | Uterine carcinosarcoma |
| UVM | Uveal melanoma |

**Figure S1.**
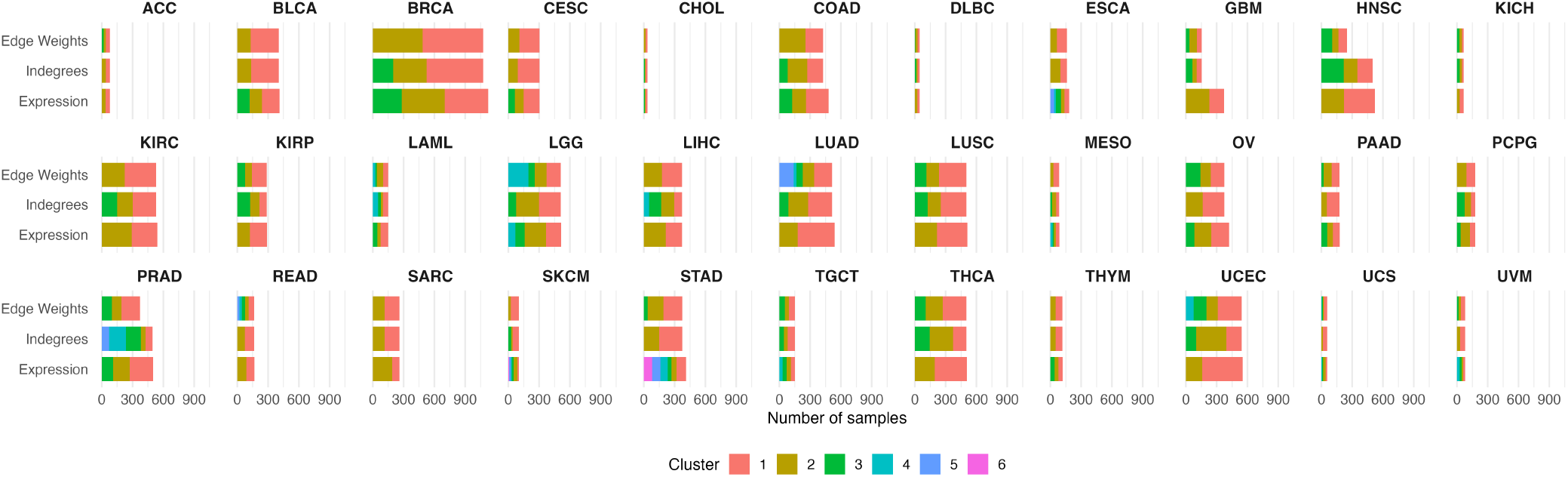
Sample distribution across clusters for the 33 TCGA cancer types.

**Figure S2.**
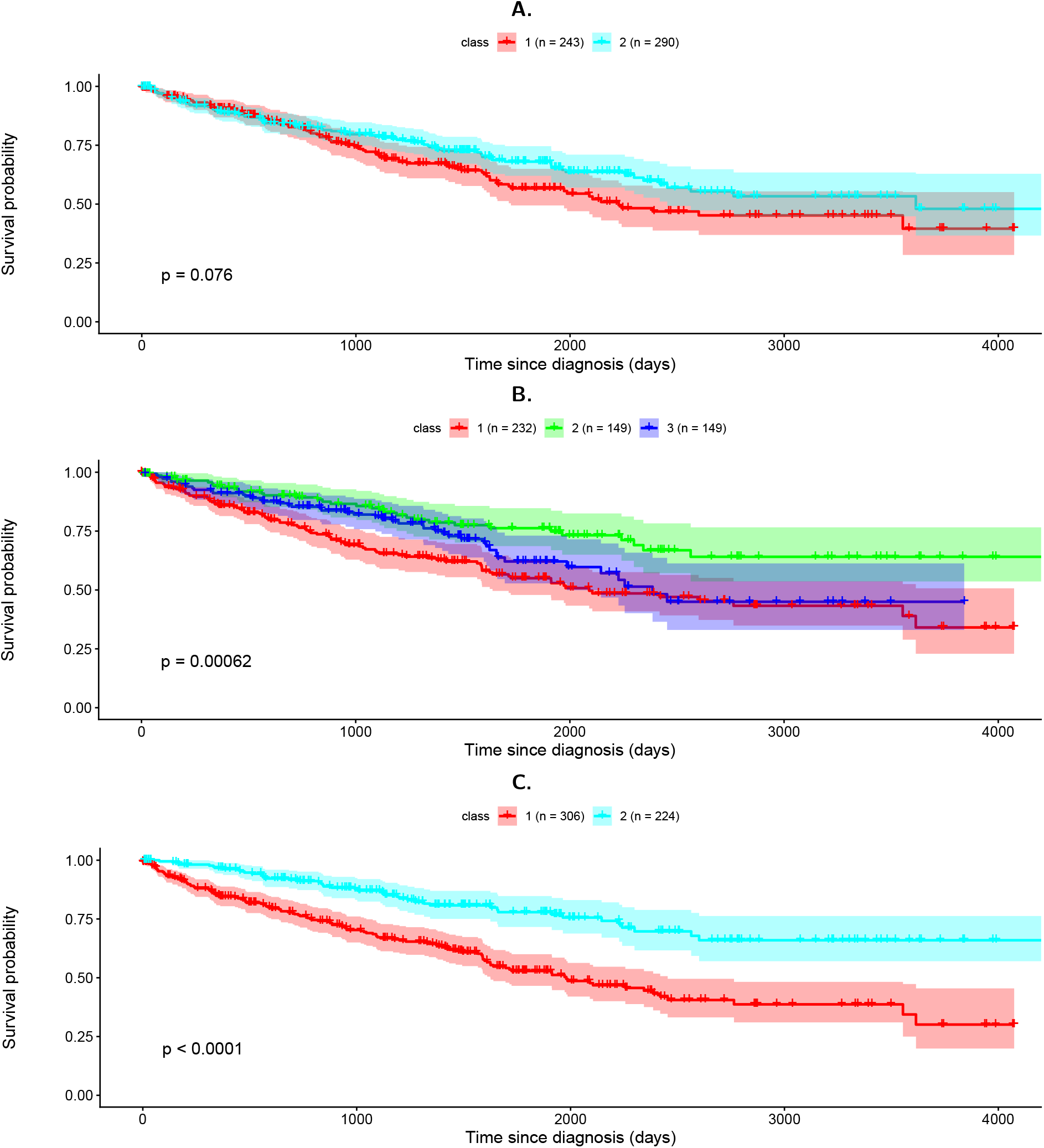
Kaplan–Meier survival analysis for KIRC. Panels A–C show survival curves for expression-based, indegree-based, and edge-weight-based clusters, respectively.

**Table S2.** Comparison of methods considered for patient-level clustering of gene expression and GRN-derived feature matrices.

| Package / Method | Language | Input Type | Relevance to This Study | Limitations for This Analysis |
| --- | --- | --- | --- | --- |
| cola (chosen) | R | Numeric feature matrix with samples as columns, including gene expression or GRN-derived matrices | Performs consensus clustering with feature selection, multiple clustering algorithms, PAC, silhouette, best-*k* selection, and automated reporting | Computationally intensive for very high-dimensional edge-weight matrices |
| ConsensusClusterPlus | R | Numeric feature matrix | Performs consensus clustering of samples using resampling-based stability assessment | Does not include built-in feature selection or the same range of diagnostics and best-*k* guidance; performed less well on our data |
| diceR | R | Numeric feature matrix | Provides ensemble clustering and can combine results from multiple clustering algorithms | Does not provide the same integrated feature-selection workflow, diagnostics, or optimal-solution guidance required for our benchmark |
| PINSPlus | R | Numeric matrix or multi-omics data | Perturbation-based sample clustering, including applications to multi-omics datasets | Does not provide the same feature-selection and diagnostic framework; produced unstable clusters with low silhouette values on our data |
| ClustOmics | Python | Multi-omics matrices or existing clustering outputs | Designed for integrating clustering information across omics layers | Not designed as a standalone method for benchmarking feature selection and clustering across expression, indegree, and edge-weight matrices |

*Note:* Graph community-detection approaches, including Infomap, MCL, CDlib, fastconsensus, and netneurotools, were also considered but were not included in the final comparison because they identify communities of nodes within a single graph or adjacency matrix, whereas our objective was to cluster patients across a collection of patient-specific GRNs. WGCNA was similarly excluded because it identifies gene co-expression modules rather than unsupervised patient subgroups.

**Figure S3.**
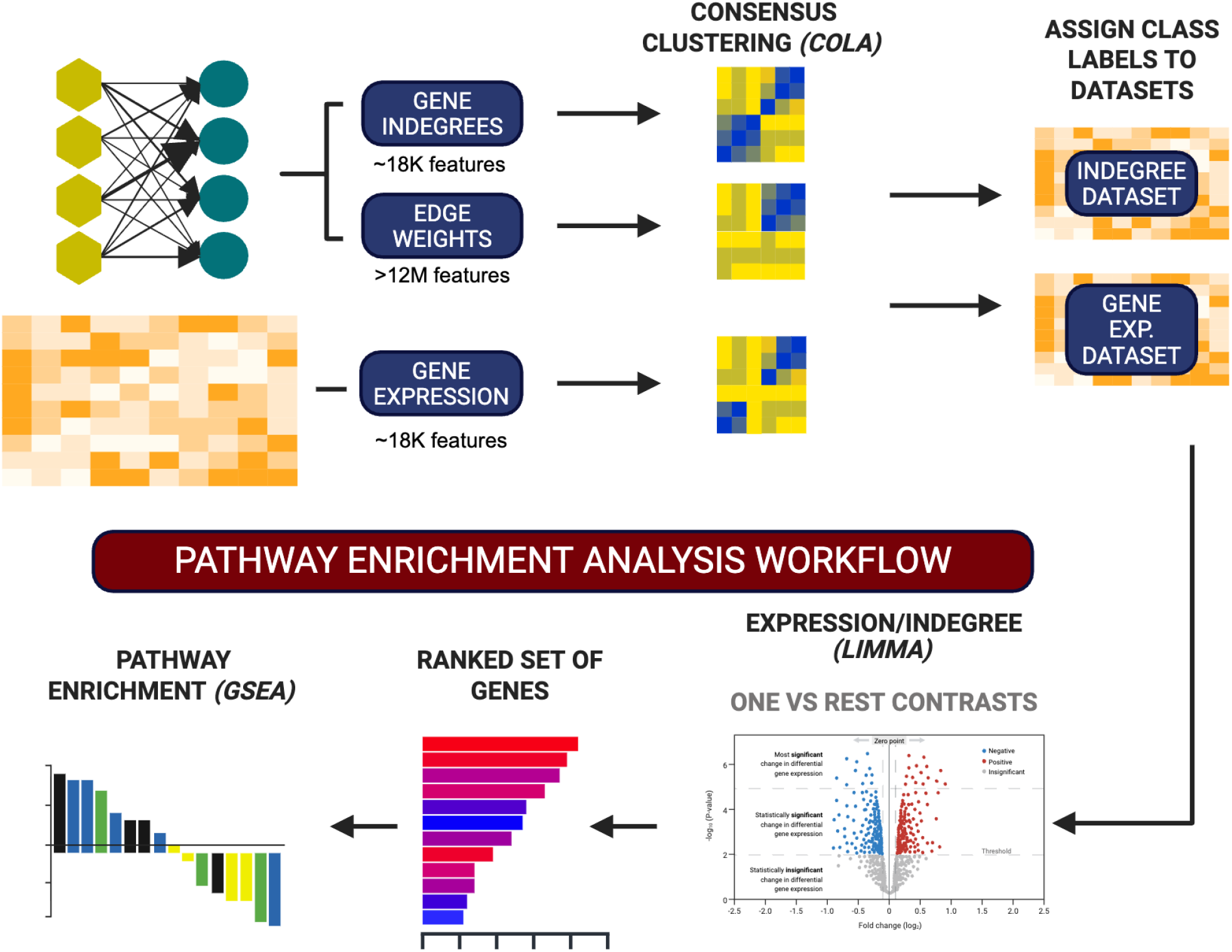
Workflow for pathway enrichment analysis of clusters derived from indegree, edge-weight, and gene expression features.

**Figure S4.**
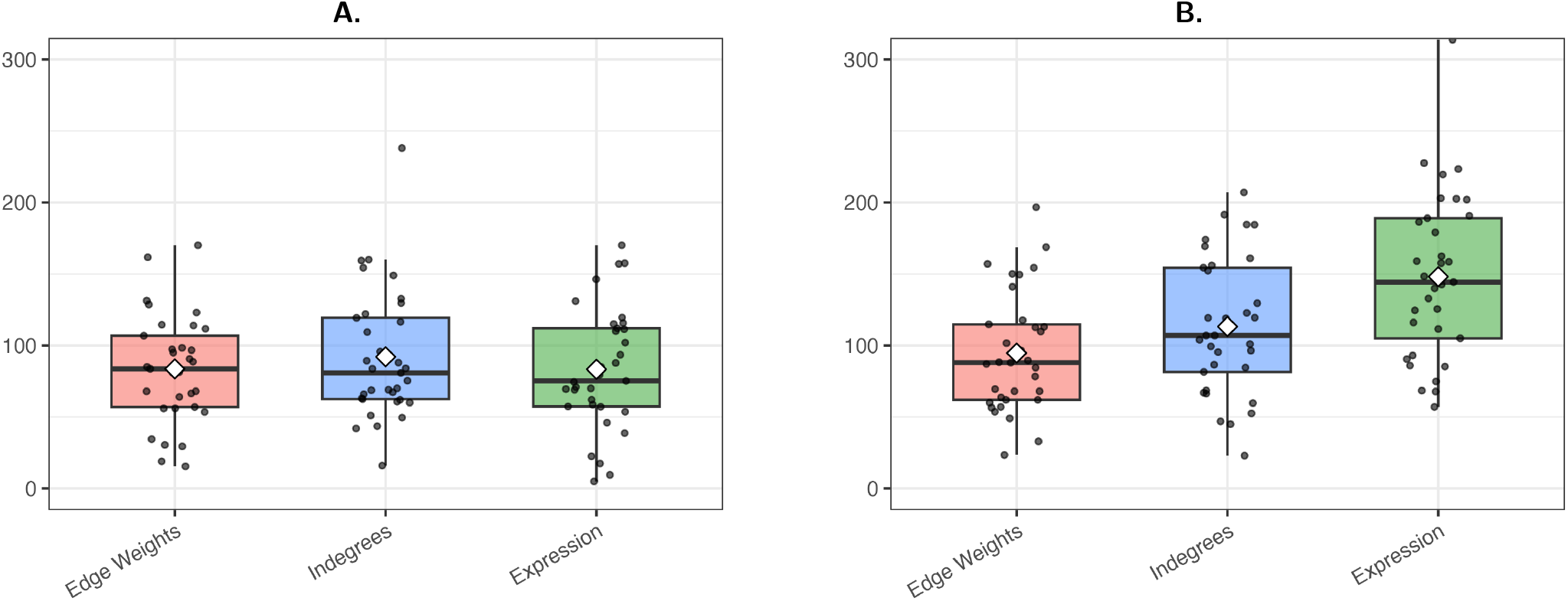
Summary of the number of unique Reactome pathways identified by GSEA across clustering modalities. Boxes indicate the interquartile range, horizontal lines indicate medians, and white diamonds indicate mean values. (A) Average number of unique pathways identified when genes were ranked using indegree values, with clusters defined from each data type. (B) Average number of unique pathways identified when genes were ranked using gene expression values, with clusters defined from each data type.

**Table S3.**
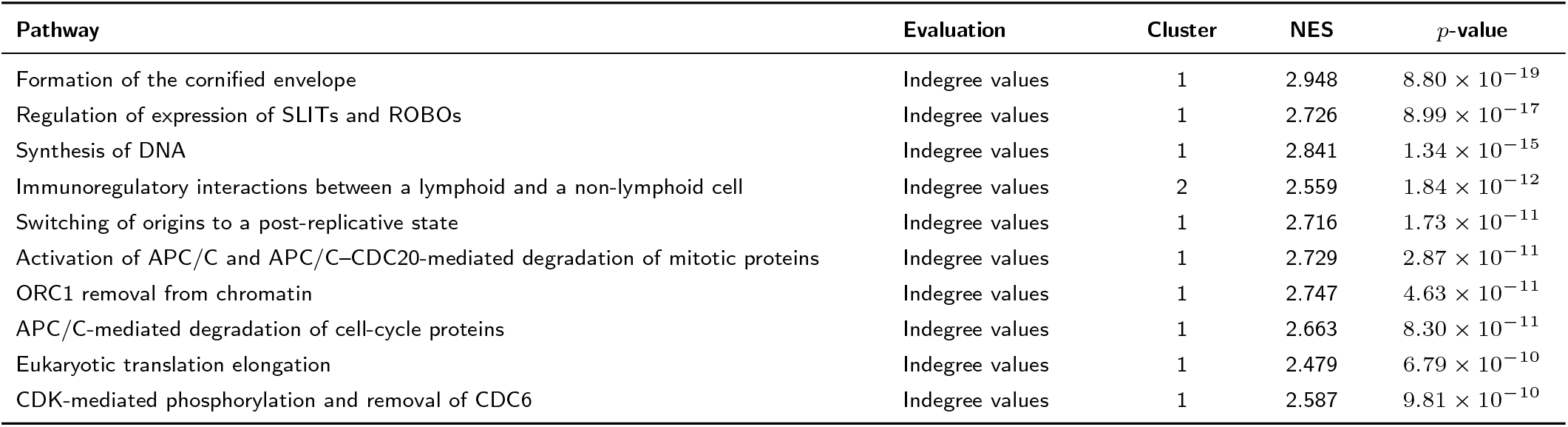
Reactome pathway enrichment results for the ten pathways with the lowest *p*-values identified from edge-weight-derived KIRC clusters, considering GSEA performed using either gene expression or indegree values. The table reports the cluster in which each pathway showed positive enrichment, the normalized enrichment score (NES), and the corresponding *p*-value.

| Pathway | Evaluation | Cluster | NES | $p$-value |
| --- | --- | --- | --- | --- |
| Formation of the cornified envelope | Indegree values | 1 | 2.948 | $8.80 \times 10^{-19}$ |
| Regulation of expression of SLITs and ROBOs | Indegree values | 1 | 2.726 | $8.99 \times 10^{-17}$ |
| Synthesis of DNA | Indegree values | 1 | 2.841 | $1.34 \times 10^{-15}$ |
| Immunoregulatory interactions between a lymphoid and a non-lymphoid cell | Indegree values | 2 | 2.559 | $1.84 \times 10^{-12}$ |
| Switching of origins to a post-replicative state | Indegree values | 1 | 2.716 | $1.73 \times 10^{-11}$ |
| Activation of APC/C and APC/C–CDC20-mediated degradation of mitotic proteins | Indegree values | 1 | 2.729 | $2.87 \times 10^{-11}$ |
| ORC1 removal from chromatin | Indegree values | 1 | 2.747 | $4.63 \times 10^{-11}$ |
| APC/C-mediated degradation of cell-cycle proteins | Indegree values | 1 | 2.663 | $8.30 \times 10^{-11}$ |
| Eukaryotic translation elongation | Indegree values | 1 | 2.479 | $6.79 \times 10^{-10}$ |
| CDK-mediated phosphorylation and removal of CDC6 | Indegree values | 1 | 2.587 | $9.81 \times 10^{-10}$ |

